# Interannual climate variability data improves niche estimates in species distribution models

**DOI:** 10.1101/2021.08.30.458152

**Authors:** Dirk Nikolaus Karger, Bianca Saladin, Rafael O. Wüest-Karpati, Catherine H. Graham, Damaris Zurell, Lidong Mo, Niklaus E. Zimmermann

## Abstract

**Aim:** Climate is an essential element of species’ niche estimates in many current ecological applications such as species distribution models (SDMs). Climate predictors are often used in the form of long-term mean values. Yet, climate can also be described as spatial or temporal variability for variables like temperature or precipitation. Such variability, spatial or temporal, offers additional insights into niche properties. Here, we test to what degree spatial variability and long-term temporal variability in temperature and precipitation improve SDM predictions globally.

**Location:** Global.

**Time period:** 1979-2013

**Major taxa studies:** Mammal, Amphibians, Reptiles

**Methods:** We use three different SDM algorithms, and a set of 833 amphibian, 779 reptile, and 2211 mammal species to quantify the effect of spatial and temporal climate variability in SDMs. All SDMs were cross-validated and accessed for their performance using the Area under the Curve (AUC) and the True Skill Statistic (TSS).

**Results:** Mean performance of SDMs with climatic means as predictors was TSS=0.71 and AUC=0.90. The inclusion of spatial variability offers a significant gain in SDM performance (mean TSS=0.74, mean AUC=0.92), as does the inclusion of temporal variability (mean TSS=0.80, mean AUC=0.94). Including both spatial and temporal variability in SDMs shows similarly high TSS and AUC scores.

**Main conclusions:** Accounting for temporal rather than spatial variability in climate improved the SDM prediction especially in exotherm groups such as amphibians and reptiles, while for endotermic mammals no such improvement was observed. These results indicate that more detailed information about temporal climate variability offers a highly promising avenue for improving niche estimates and calls for a new set of standard bioclimatic predictors in SDM research.

## Introduction

Climate is known to influence species’ distributions (Woodward & Woodward, 1987) across different spatial and temporal scales, thus, detecting the impact of climate on species depends to a large degree on the spatial and temporal scale at which it is assessed (Wiens, 1989; Rahbek, 2005). While the effect of spatial variability in climate on species distributions has received heightened attention in recent years due to the ever increasing spatial resolution of freely available climate data (e.g. Hijmans *et al*., 2005; Fick & Hijmans, 2017; Karger *et al*., 2017), analyses of the effect of temporal climate variability on species ranges have been much less abundant (Zimmermann *et al*., 2009). Analytically, an increase in resolution in one dimension often comes at the cost of decreased resolution in the other dimension due to computational limitations (Hourdin *et al*., 2017; Schär *et al*., 2019). However, a focus on ever increasing spatial resolution may leave time-dependent phenomena undetected (Wiens, 1989). This is especially of concern since one of the major components of climate change is an increase in climate variability and – in consequence – an increase of extremes (Rahmstorf & Coumou, 2011; Seneviratne *et al*., 2012). Increasing frequencies and severities of extreme events may cause greater physiological stress and may thus result in rapid responses in many species with likely severe consequences for their spatial distribution (Reyer *et al*., 2013; Alexander *et al*., 2015).

The spatial ranges of species are commonly estimated using empirical species distribution models (SDMs), sometimes also termed bioclimatic envelope or habitat suitability models (Guisan & Zimmermann, 2000; Guisan & Thuiller, 2005). SDMs characterize the environmental niche of a species (Hutchinson, 1957) usually with respect to a few key environmental factors, such as temperature and precipitation. It has become common practice to use a limited set of climate variables as predictor variables (Araújo & Guisan, 2006) based on long-term means (climatologies) alone (Ashcroft *et al*., 2011). Such an aggregation of climate variability into long-term climatological means fully removes information on the temporal signal, including inter-annual variability. Species are known to strongly differ in degree to which they can tolerate climatic extremes, and this affects their life cycles, coping strategies through functional adaptations and, ultimately, their spatial distribution. Yet when using long term climatic means, our capacity to distinguish effects from differences in climate variability on species’ distributions is basically removed.

Climatologies do not only smooth out temporal variabilities in climate but also reduce spatial variability. Spatial climate heterogeneity as a result of small-scale topography and other factors are not represented in gridded datasets of coarse spatial grain. Coarse spatial grains cannot resolve the richness in topography, and thus climate, environment, and habitats, that are essential for quantifying species’ environmental niches (Stein *et al*., 2014; Stein & Kreft, 2015). Such misrepresentation of spatial heterogeneity when representing or aggregating climate predictors at coarse grain might more strongly impact niche estimates in the tropics compared to temperate or boreal zones (Janzen, 1967). In the tropics, species generally experience a lower degree of intra- and interannual climatic variation due to the rather stable environmental conditions they encounter throughout the year (Janzen, 1967; Wiens, 1989). In temperate climates however, the conditions a species experiences are much more variable due to the larger intra- and interannual variation in climate (Janzen, 1967). Species occurring in tropical ecosystems, therefore often have a much narrower climatic niche (Stevens, 1989; Cadena *et al*., 2012). In turn, this also implies that the influence of temporal variability might be greater in areas where species are not well adapted to variation in climate. Therefore, along large-scale geographic gradients both spatial and temporal variability can become important in estimations of a species environmental niche.

Climate does not only influence the distribution of species, but also has a profound impact on how specific traits evolve over time as adaptations to climate itself (Kozak & Wiens, 2010; Rolland *et al*., 2018; Liu *et al*., 2020). While climate is an important factor in the diversification of species (Liu *et al*., 2020), many adaptations can also be directly linked to environmental factors. A prominent example is the evolution of endothermy, which allows to some degree, the decoupling of a species internal temperature from that of its surrounding habitat (McNab, 1978; Ruben, 1995), which in turn leads to different responses of species to a climatic factor. Hence, overall, spatial and temporal variabilities may not only act as species range determinants in isolation, but also interact with each other. With climatic data becoming available at increasingly higher spatial and temporal resolution, the opportunity arises to generate an improved understanding of the role of spatial and temporal climatic variability on the distribution of species. Here, we evaluate if considering temporal, and spatial variation in addition to classical coarse-grained climate mean values improves the performance of SDMs when using coarse-grained species distribution data.

Based on the potential effects of spatial and temporal climate variability on species discussed before, we hypothesize that

i. both the inclusion of spatial and temporal variability positively affect the performance of SDMs,
ii. the performance of ectotherm SDMs increases more strongly when accounting for spatial and temporal variability than the performance of endotherm SDMs, and
iii. SDMs for tropical and mountain species will benefit more strongly from including variability than SDMs of species from other habitats.

We test these hypotheses by modelling the distribution of 833 amphibian, 779 reptile, and 2211 mammal species as a function of current climate using four different predictor groups composed of different combinations of input variables: mean climate, spatial climatic variability and temporal (interannual) climatic variability.

## Methods

### Species data

We used global distribution maps provided by the Amphibian, Mammal, and Reptile Red List Assessment (IUCN, 2016). Grid cells within the distribution range of each species were converted to 0.5° grid cells, which is close to 50 km at the equator, a resolution suggested as appropriate (Hurlbert & Jetz, 2007) and often used (Fritz & Rahbek, 2012; Zurell *et al*., 2018; Thuiller *et al*., 2019) when gridding polygon range maps at the global scale. Grid cells intersecting with a range map polygon were assigned as presence cells, while those not intersecting where treated as absence cells. We only considered species for which the presences cover at least 72 0.5° grid cells so that a minimum of six data points per predictor variable (including quadratic terms) was available for model building. We also removed domestic and aquatic species.

### Climate predictor groups

We used global climate data from CHELSA V1.2 (Karger *et al*., 2017a,b) and built several groups of predictors (Fig. 1, Table 1) by aggregating CHELSA to the 0.5° grid of the species data by taking the mean of all 30 arc second grid cells overlapping with a 0.5° grid cell. To calculate sub-grid heterogeneity of a climatic variable (hereafter: spatial) within a 0.5° grid cell, we used the standard deviation of all CHELSA 30 arc second grid cells overlapping with 0.5° grid cells. To calculate the interannual variability (hereafter: temporal) we calculated the standard deviation of mean annual 2m air temperature for each year from 1979 to 2013 from CHELSA V1.2 per grid cell. For temporal precipitation variability we used the relative standard deviation (temporal RSD, equivalent to the coefficient of variation) of the annual precipitation sum across all years from 1979 to 2013 from CHELSA V1.2 per grid cell. Based on the data aggregated as explained above, we generated four different groups of predictors for annual temperature and precipitation, with different combinations of spatial and temporal variabilities (Table 1).

**Fig. 1.**
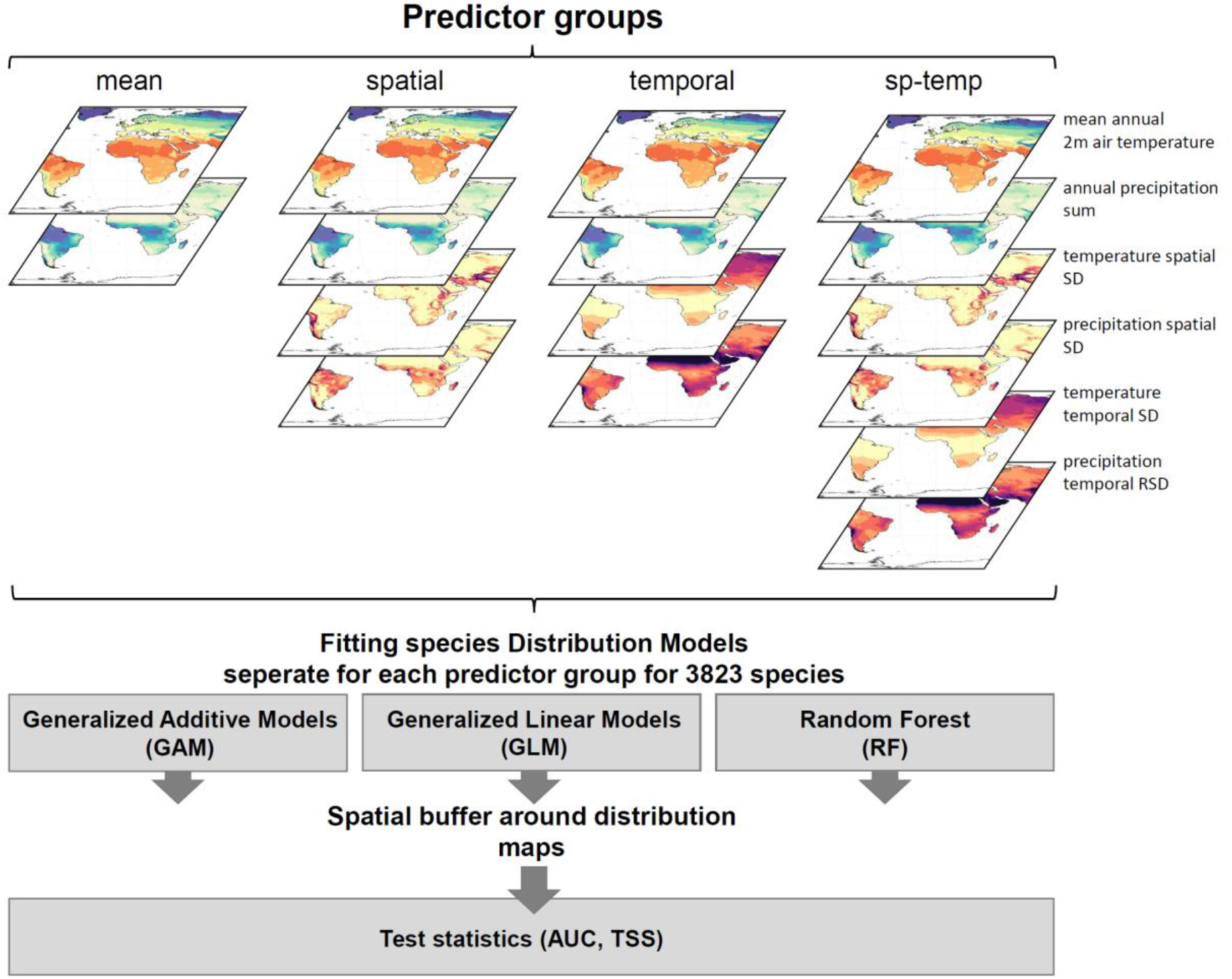
Schematic representation of the analytical setup. Four predictor groups where formed and three algorithms for species distribution models (SDMs) were fitted from range maps for 3823 species of mammals, amphibians, and reptiles. The different SDMs were predicted spatially and their predictive performance assessed within a buffer of 3000 km around observed ranges, using the area under the curve (AUC) and the true skill statistic (TSS) as performance measures.

**Table 1.** Characterization of the four predictor groups used with the respective variables included in each group as well as the temporal or spatial unit over which they were aggregated. All variables are based on CHELSA V1.2 (Karger *et al*., 2017a).

| Predictor group | variables included | aggregation unit |
| --- | --- | --- |
| mean | mean annual 2m air temperature 1979-2013 | - |
|  | mean annual precipitation sum 1979-2013 | - |
| spatial | mean annual 2m air temperature 1979-2013 | - |
|  | mean annual precipitation sum 1979-2013 | - |
|  | standard deviation mean annual 2m air temperature 1979-2013 | all 0.0083334° grid cells within a 0.5° grid cell |
|  | standard deviation mean annual precipitation sum 1979-2013 | all 0.0083334° grid cells within a 0.5° grid cell |
| temporal | mean annual 2m air temperature 1979-2013 | - |
|  | mean annual precipitation sum 1979-2013 | - |
|  | standard deviation mean annual 2m air temperature 1979-2013 | all years from 1979-2013 |
|  | coefficient of variation mean annual precipitation sum 1979-2013 | all years from 1979-2013 |
| sp-temp | mean annual 2m air temperature 1979-2013 | - |
|  | mean annual precipitation sum 1979-2013 | - |
|  | standard deviation mean annual 2m air temperature 1979-2013 | all 0.0083334° grid cells within a 0.5° grid cell |
|  | standard deviation mean annual precipitation sum 1979-2013 | all 0.0083334° grid cells within a 0.5° grid cell |
|  | standard deviation mean annual 2m air temperature 1979-2013 | all years from 1979-2013 |
|  | coefficient of variation mean annual precipitation sum 1979-2013 | all years from 1979-2013 |

### Species distribution modelling

We used three algorithms to relate presences and absences with the selected environmental predictor sets: Generalized Linear models (GLM) (Nelder & Wedderburn, 1972), Genearlized Additive Models (GAM) (Hastie & Tibshirani, 1990), and Random Forests (RF) (Breiman, 2001). GLMs were run using linear and quadratic terms, GAMs was run using thin plate splines setting an upper limit of 4 degrees of freedom (k=5). In both cases, we set weights such that the sum of weights of presences equaled the sum of weights of absences (Barbet‐Massin *et al*., 2012). A classification RF was fitted using 1500 trees, while sub-sampling was restricted to contain equal numbers of presences and absences.

To assess model performance, we tested SDM predictions only within a buffer around each species’ range polygon. By doing so we account for biogeographic history and explicitly test how well a model predicts the actual range of a species rather than how well it also makes predictions far outside a species’ range, yet with suitable climate. We applied a buffer of 3000km around each range polygon and fitted and tested SDMs only within this extent.

We evaluated the predictive performance of the SDMs using repeated split-sample tests: we split the data repeatedly into 80% training and 20% test data, fitted the model on the training data, and predicted it to the test data. This procedure was repeated 30 times, while we recorded predictive performance of each repeat. Predictive performance was assessed using a) the true skills statistic (TSS) (Allouche *et al*., 2006), after thresholding the predictions into presence/absence using a TSS-optimized threshold, and b) the area under the curve (AUC) (Swets, 1988). We provide the full SDM description following the ODMAP protocol (Zurell *et al*., 2020)(Zurell et al. 2020) in Appendix S2.

### Performance tests of predictor groups

We used a linear mixed effects model (Bates *et al*., 2015) with either TSS or AUC as response variable and the predictor group as fixed effects together with the SDM algorithm (GLM, GAM, RF) and the species identity as random effects. Adding the type of SDM (GLM, GAM, or RF) as random effect on the intercept considers that algorithms can perform differently well (e.g. have a different mean performance between AUC or TSS, Thuiller *et al*., 2019). To always compare the same species, but modeled with different sets of predictor groups, we also added the identity of the species as random affect to the intercept. To check if there are differences in SDM performance across climatologies, we ran a paired Wilcoxon test. By this, we tested if one climatology performs better then another.

All analyses have been performed using the R language for statistical computing (R Core Team, 2015, version 3.6.1), the R packages raster (Hijmans & van Etten, 2014), mgcv (Wood & Wood, 2015), and randomForest (randomForest: Breiman and Cutler’s Random Forests for Classification and Regression).

### Spatial performance of different predictor groups

To test if different predictor groups have different performances in different regions, we used the gridded range map at 0.5° resolution from IUCN and assigned the value of the respective test metric (TSS, AUC) to the entire range in which a species is present. All ranges were then stacked and the mean of all TSS and AUC values covering a 0.5° grid cell calculated.

## Results

### Spatial patterns of the predictor groups

While mean annual 2m air temperatures and annual precipitation sums were generally higher in the tropics and decrease towards the poles (Fig. 2, upper), the spatial SD of these two variables is usually highest in mountainous terrain (Fig. 2, middle). Interannual variability (temporal SD) of temperature is generally higher in the northern hemisphere compared to the southern hemisphere, and increases from tropics to continental artic or boreal areas. Variability (temporal RSD) of precipitation is more idiosyncratic, with lowest values estimated in desert areas.

**Fig. 2.**
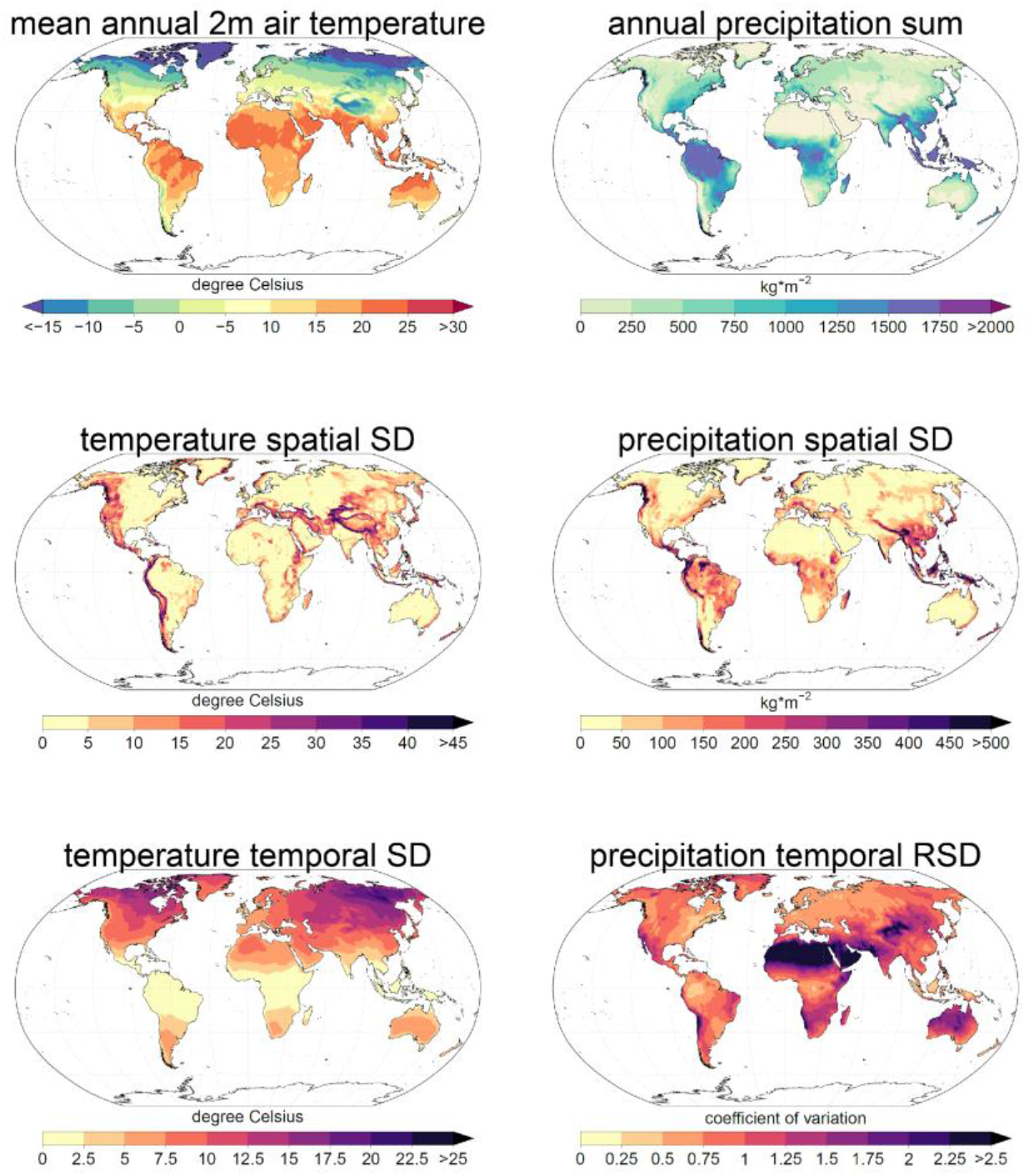
Spatio-temporal variation in 2m air temperature and precipitation used as predictors in the different SDMs based on CHELSA V1.2. Mean annual values (upper row) show the annual mean for temperature, and the mean annual sum for precipitation averaged over the years 1979-2013 and aggregated to 0.5° from a 30 arc second spatial grain by taking the mean of a 0.5° grid cell. Spatial variation (middle) indicates the standard deviation (SD) of all temperature values of a 30 arc second grid of temperature or precipitation overlapping with a 0.5° grid cell. Temporal variation (lower row) shows the standard deviation (SD) of temperature over years between 1979-2013, and the relative standard deviation (RSD) calculated as the coefficient of variation for precipitation over the same time period.

### Performance scores of the SDMs with different predictor groups

Overall the predictive performance of the SDMs was high with an average AUC of 0.92 and ranging on from 0.90 to 0.95 between different groups of predictors, SDMs, and climatologies. For TSS, values averaged 0.75 with a minimum 0.68 and a maximum of 0.82 for the different predictor groups.

SDMs based on mean climate predictors performed worst among all groups, with average AUC and TSS scores of 0.90 and 0.71, respectively (GAM: 0.91; 0.73, GLM: 0.90; 0.73, RF: 0.90; 0.68). SDMs based on mean climate predictors plus spatial predictors performed slightly better, with average AUC and TSS scores of 0.92 and 0.74 (GAM: 0.93; 0.77, GLM: 0.92; 0.76, RF: 0.91; 0.70). SDMs based on mean climate predictors plus temporal predictors performed slightly better, with average AUC and TSS scores of 0.94 and 0.80 (GAM: 0.94; 0.81, GLM: 0.94; 0.81, RF: 0.94; 0.77). SDMs containing mean climate predictors plus spatial and temporal predictors performed similar as the SDMs with mean and temporal predictors, with average AUC and TSS scores of 0.94 and 0.80 (GAM: 0.95; 0.82, GLM: 0.93; 0.81, RF: 0.94; 0.77). The results of the linear mixed effects model to assess the SDM performance with different predictor groups showed that adding predictors that account for either spatial or temporal variation increased predictive performance of models across all groups of vertebrates. Models based on the temporal predictor group outperformed models based on the spatial predictor group for amphibians and reptiles, but not for mammals (equal performance). Models that included predictors that accounted for both spatial and temporal variation (sp-temp predictor group) performed best across all vertebrate groups. Fig. 3 illustrates these results using effect-size plots (below boxplots) for TSS, while results for AUC are equivalent (see Supplemental Figure S1).

**Fig. 3.**
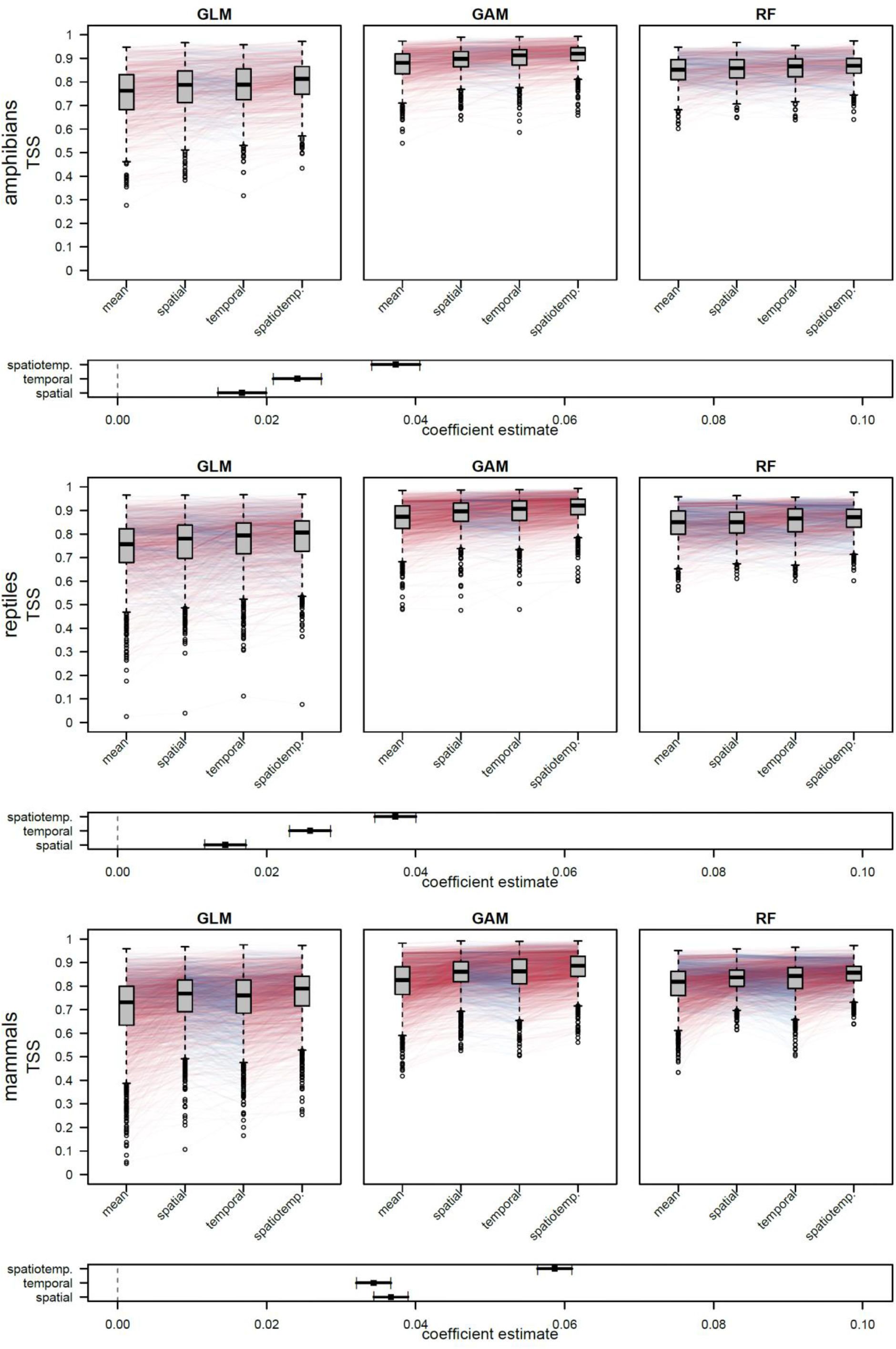
Comparison of the performance of three different SDM algorithms (GLM = Generalized linear model, GAM = Generalized additive model, RF = Random Forests) calculated with four different sets of predictors for amphibians, reptiles, and mammals measured by the True Skill Statistic (TSS). Colored lines connect pairs of SDMs based on different predictor sets for the same species, with red and blue lines indicating pairs in which TSS values increased and decreased between predictor groups from left to right. Plots below the boxplots shows the coefficient estimates of a linear mixed effects model with TSS as response, the groups (mean, spatial, temporal, sp-temp) as predictor, and the model (GLM, GAM, RF) as well as the species ID as random effects. Coefficients are in relation to the performance of SDMs with the predictor set: mean.

### Spatial comparison

The performance of SDMs is highly variable across the globe (Fig. 4). The mean predictor group generally performed worst in mountainous terrain, such as the Andes or the Himalayas, but also Madagascar showed low TSS and AUC scores (Fig. 4, Supplementary Fig. S2 for AUC). Including spatial variability in temperature and precipitation in SDMs improved the models in these areas, but showed a slight decline in performance in desert and arctic areas (Fig. 4: TSS differences spatial-mean). Adding temporal variability to the models containing mean predictors resulted in improved SDM performance in almost all areas (Fig. 4: temporal-mean). Including all spatial and temporal predictors resulted in a slight improvement in the spatial model accuracies compared to including the temporal group with the mean predictor group (Fig. 4: spatio temporal), yet the improvement compared to mean plus temporal SDM were small.

**Fig 4.**
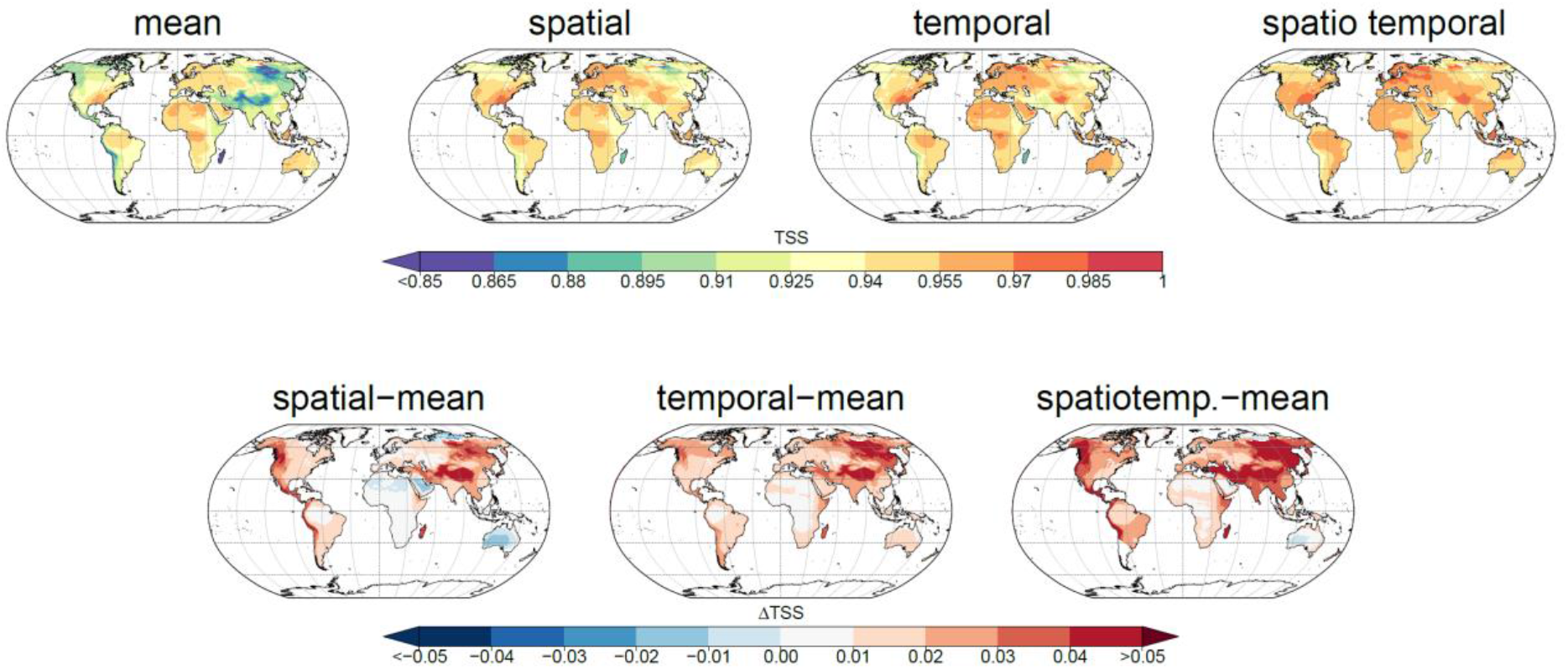
Spatial variation in mean TSS values per grid cell and TSS differences between models using different predictor groups. The upper row illustrates TSS averaged for all mammals, reptiles, and amphibians modeled for the four models using different predictor groups. The lower row illustrates the averaged TSS difference among all SDMs when adding either spatial, temporal or both spatial and temporal (spatiotemp.) predictors to SDMs based on mean predictors only.

## Discussion

Including temporal variability of predictors into broad-scale SDMs leads to a greater improvement of model performance than the inclusion of sub-grain spatial variability of these predictors. These findings suggest that especially the inclusion of interannual climate variability has a large potential of improving the estimation of niche characteristics across a large range of taxa. Including temporal predictor variability increased the performance of SDMs in almost all areas across the globe, though to differing degrees. Especially prominent is the increase in areas with marked seasonality such as in tropical monsoon climates, tropical wet and dry climate, or areas that receive very infrequent precipitation such as the Horn of Africa (Beck *et al*., 2018). Temporal variability does also increase the performance of SDMs in mountainous regions potentially indicating that temporal variability in a climate variable is also capable to capture the niche limitations that are otherwise captured by spatial heterogeneity. One reason for the performance gain when including temporal variability is that it expresses the degree of climatic extremes which can physiologically limit the distribution of species (Zimmermann *et al*., 2009). Although, the degree to which extremes are represented in such variability predictors certainly depends on the temporal resolution of the climatic input dataset. In the case presented here, we used interannual variation, which means that extreme events are restricted to extremely dry or wet years, or extremely hot or cold years. Using more detailed temporal analyses would allow to refine the representation of climatic extremes further.

As expected, the inclusion of the spatial variability improves SDMs mainly in mountainous areas where climate is extremely heterogeneous over short distances. The improvement was specifically strong in tropical mountains where species usually occur in narrow elevational bands with little or no intra-annual variability (Janzen, 1967; Ghalambor *et al*., 2006). In topographically less heterogenous terrain however, we observed a decline in the predictive power of SDMs. Almost all over Africa, Australia, and the low elevation parts of Eurasia, and North America spatial variability has no effect, or even a negative effect on the performance of SDMs. In these areas spatial heterogeneity is low and inclusion of spatial variability in climate predictors seems biologically unimportant, which leads to a decrease in their performance (Loehle & LeBlanc, 1996; Davis *et al*., 1998; Vaughan & Ormerod, 2003; Dormann *et al*., 2012).

The increase in SDM performance is however not equal across the three taxonomic groups analyzed here. While amphibians and reptiles show a significantly higher SDM performance of the temporal predictor group over the spatial predictor group, mammal SDMs do not significantly differ when either spatial or temporal predictors are added to the mean predictor group. This difference might be explained by the differences in physiology between these groups. All three groups have evolved differently in response to their environment, with ectothermic groups being much less adaptable to climatic variations than endothermic groups (Rolland *et al*., 2018). Such evolved differences in physiology ultimately affect how organisms interact with and are constrained by their environment (Buckley *et al*., 2012). Ectothermic species for example cannot buffer climate variation as well as endothermic species (Clusella-Trullas *et al*., 2011; Sunday *et al*., 2011; Hoffmann *et al*., 2013; Gunderson & Stillman, 2015) which have evolved the physiological capacity to regulate temperatures to some extent (Pither, 2003). When building SDMs from climate means alone, thus neglecting the temporal dimension of predictors, we miss out on important climatic constraint especially for endothermic species distributions, ultimately limiting the accuracy of niche estimations (Zimmermann *et al*., 2009). Model formulation and parametrization certainly plays a role in the observed differences between predictor groups. More predictors in a model usually lead to a better overall fit of a model (Brun *et al*., 2019) which can partly explain the increase in predictive power when the predictors based on mean climate are complemented with either spatial or temporal variability. As both, the spatial and the temporal predictor groups have the same number of variables, this effect does not hold when comparing these two. Combining mean with spatial and temporal predictor groups however, lead to an additional improvement. At this point however, the parametrization of has not yet plateaued (Randin *et al*., 2006; Chala *et al*., 2016; Brun *et al*., 2019; Gregr *et al*., 2019) and model performance still increases when using both spatial and temporal variability as predictors. Using different SDM algorithms mainly affects the absolute performance of the SDMs in terms of the specific test metric (AUC, TSS). However, it did not affect the relative difference in model performance between SDMs calculated from different predictor groups.

With an increasing need in biodiversity modeling for current, past, and future predictions a better understanding of the climatic predictors that quantify the ecological niche of a species is needed. Here, we show that specifically the inclusion of temporal variability offers a promising improvement in modelling the current distribution of species. Yet, also the inclusion of spatial (sub-grain) variabilities can improve model accuracies, primarily mountains and most clearly in tropical mountains. In summary, we anticipate that a more detailed inclusion of the temporal variability of climate variables offers a highly promising avenue for improving species distribution modelling in the future.

## Data Accessibility Statement

The data that support the findings of this study are openly available in EnviDat (envidat.ch) at http://doi.org/XXXXXX. Codes related to this study will be available on Zenodo at http://doi.org/YYYYYYYY

## Acknowledgements

D.N.K. & N.E.Z. acknowledge funding from: The WSL internal grants exCHELSA, as well as funding through the 2019-2020 BiodivERsA joint call for research proposals, under the BiodivClim ERA-Net COFUND program, and with the funding organisations Swiss National Science Foundation SNF (project: FeedBaCks, 193907), D.N.K., N.E.Z., & D.Z. acknowledge funding from the Swiss Data Science Projects: SPEEDMIND, and COMECO. D.N.K. & C.H.G. acknowledges funding to the ERA-Net BiodivERsA -Belmont Forum, with the national funder Swiss National Foundation (20BD21_184131), part of the 2018 Joint call BiodivERsA-Belmont Forum call (project ‘FutureWeb’). DZ also acknowledges funding from the German Science Foundation (ZU 361/1-1).

## Author contributions

D.N.K. and N.E.Z. developed the idea with input from all co-authors, D.N.K., L.M. and R.O.W. implemented the species distribution modelling, B.S. and D.N.K. analyzed the results further, D.N.K. wrote the first draft of the manuscript, and all authors contributed equally to the revisions.

## Biosketch

Dirk Nikolaus Karger is a senior researcher at the Swiss Federal Research Institute WSL and mainly interested in the impact climate has on global ecosystems and species. The author consortium binds a strong interest in species distribution modelling.

## Appendices

**Fig. S1.**
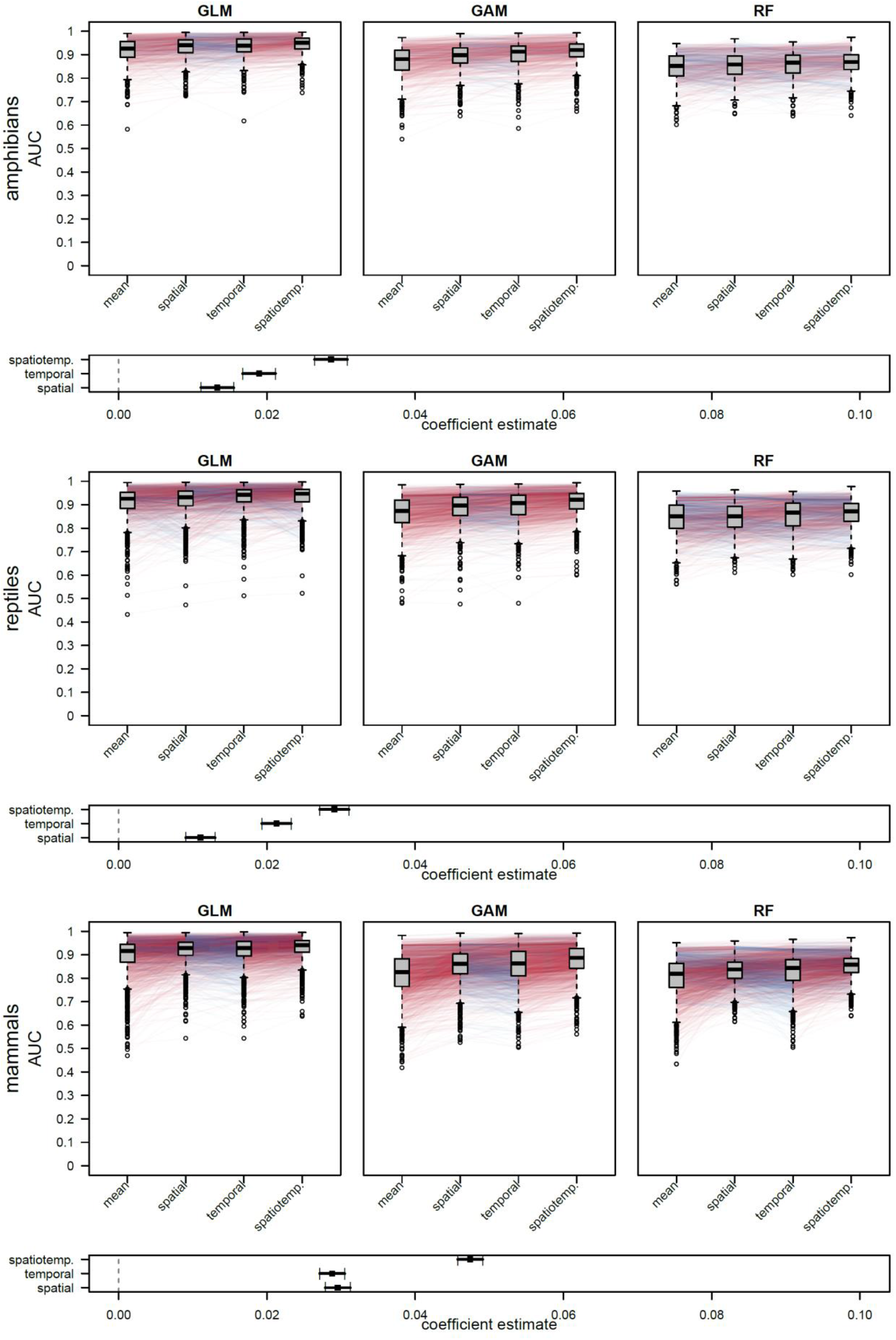
Comparison of the performance of three different SDM algorithms (GLM = Generalized linear model, GAM = Generalized additive model, RF = Random Forests) calculated with four different sets of predictors for amphibians, reptiles, and mammals measured by the Area Under the Curve (AUV). Colored lines connect pairs of SDMs based on different predictor sets for the same species, with red and blue lines indicating pairs in which AUC values increased and decreased between predictor groups from left to right. Plots below the boxplots shows the coefficient estimates of a linear mixed effects model with AUC as response, the groups (mean, spatial, temporal, spatiotemp.) as predictor, and the model (GLM, GAM, RF) as well as the species ID as random effects. Coefficients are in relation to the performance of SDMs with the predictor set: mean.

**Fig S2.**
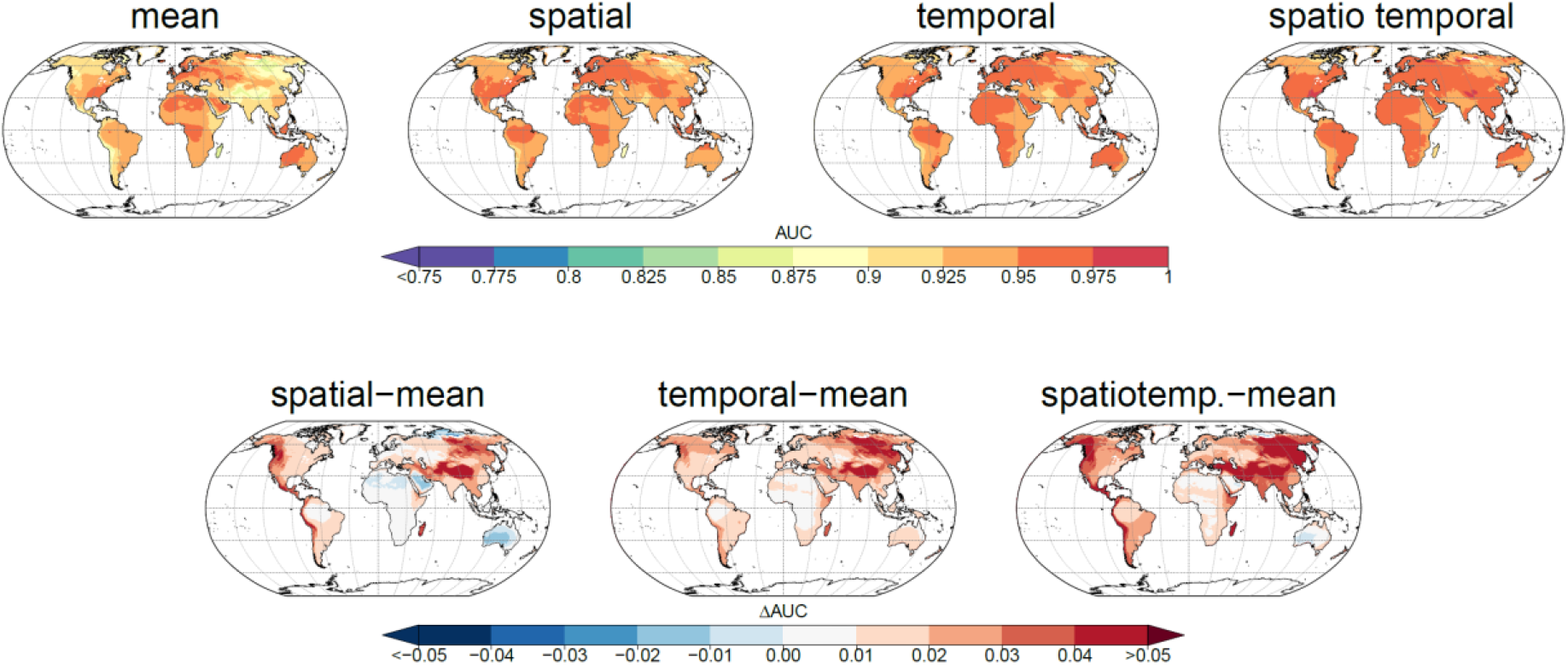
Spatial variation in mean AUC values per grid cell and AUC differences between models using different predictor groups. The upper row illustrates AUC averaged for all mammals, reptiles, and amphibians modeled for the four models using different predictor groups. The lower row illustrates the averaged AUC difference among all SDMs when adding either spatial, temporal or both spatial and temporal (spatiotemp.) predictors to SDMs based on mean predictors only.

